## Supplementary Materials for "Specialization of ubiquitin ligases to distinct nucleic acid sensors"

**The PDF file includes:**

Supplementary Figures S1 to S5

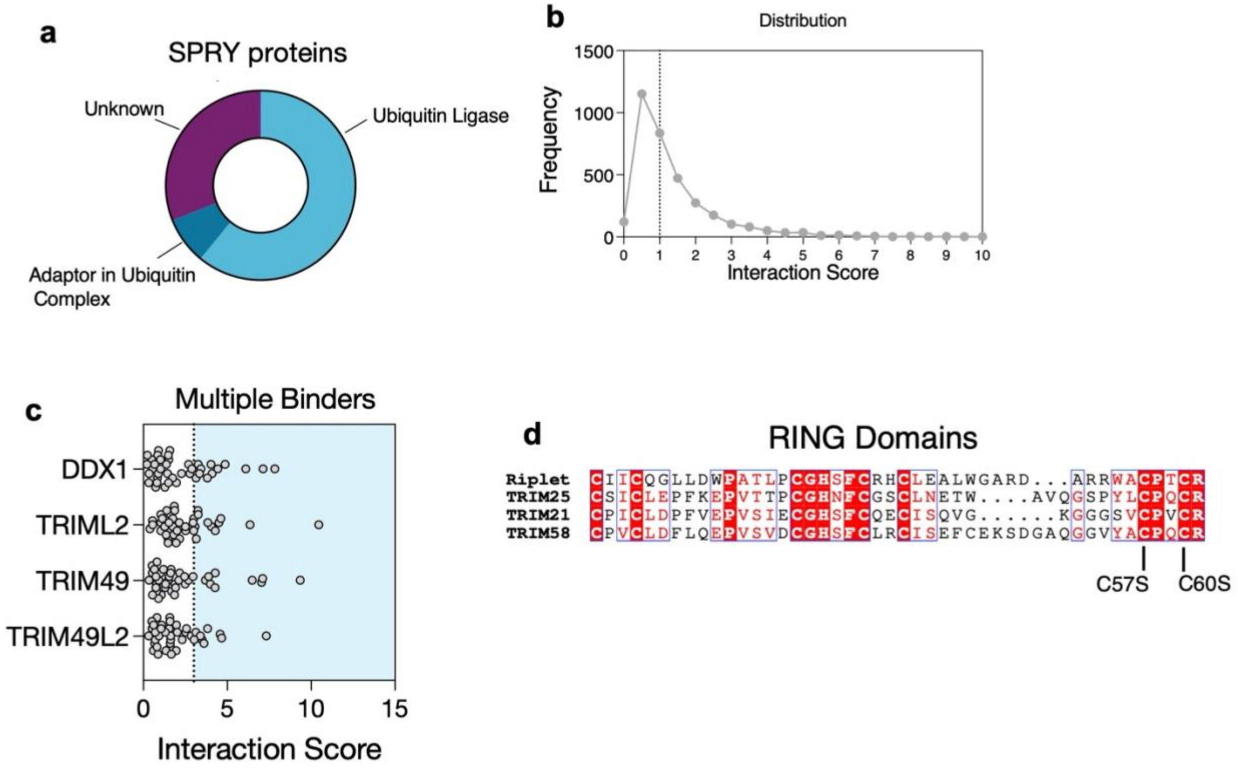


**Supplementary Figure 1. Analysis of all SPRY-Sensor predictions generated by AlphaFold**. Proportion of known biological functions of all human SPRY-containing proteins (a). Distribution of interaction scores across all SPRY-sensors pairs (b). SPRY-containing proteins with the largest number of interactor partners in the AlphaFold prediction screen (c). Multiple sequence alignment of the RING domains of Riplet, TRIM25, TRIM21 and TRIM58 (d).

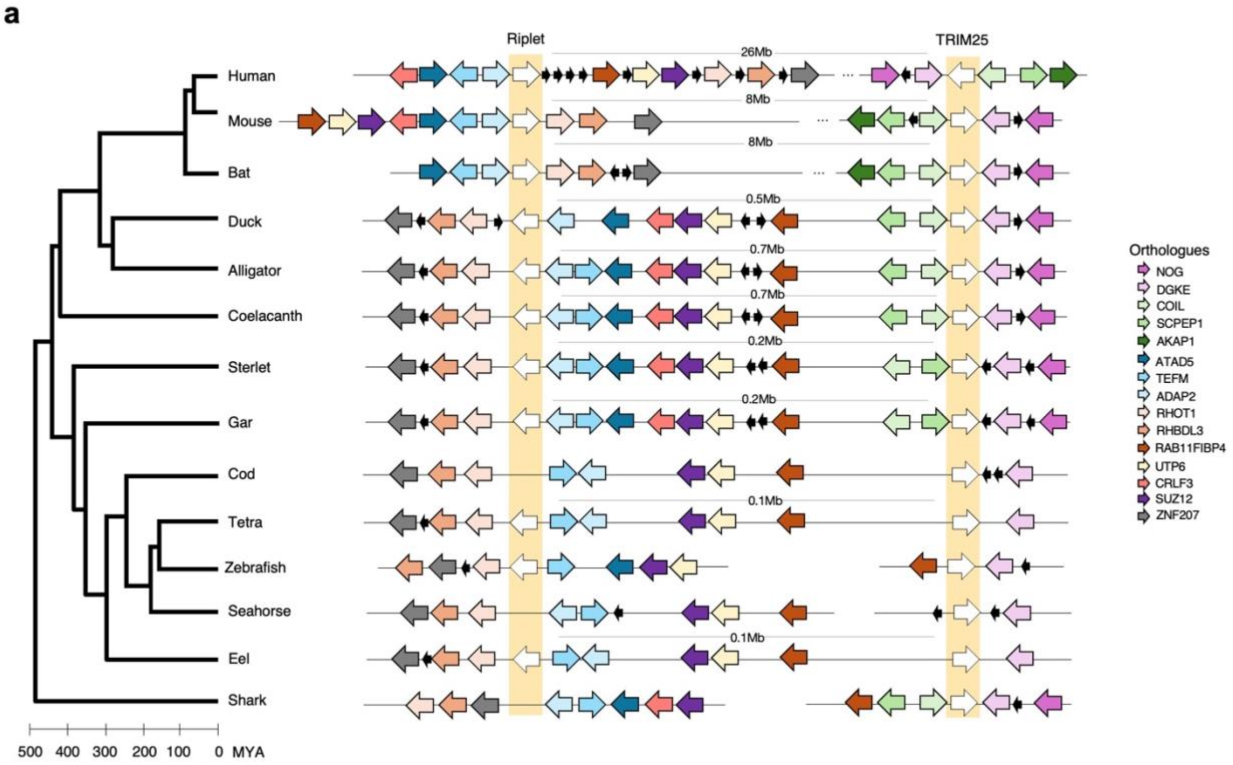


**Supplementary Figure 2. Synteny of *TRIM25* and *Riplet/RNF135* across vertebrates**. Diagram of the genetic organization of TRIM25 and Riplet loci (in yellow) in vertebrate genomes (a). Mb, megabase (1 million base pairs). MYA, Million years ago.

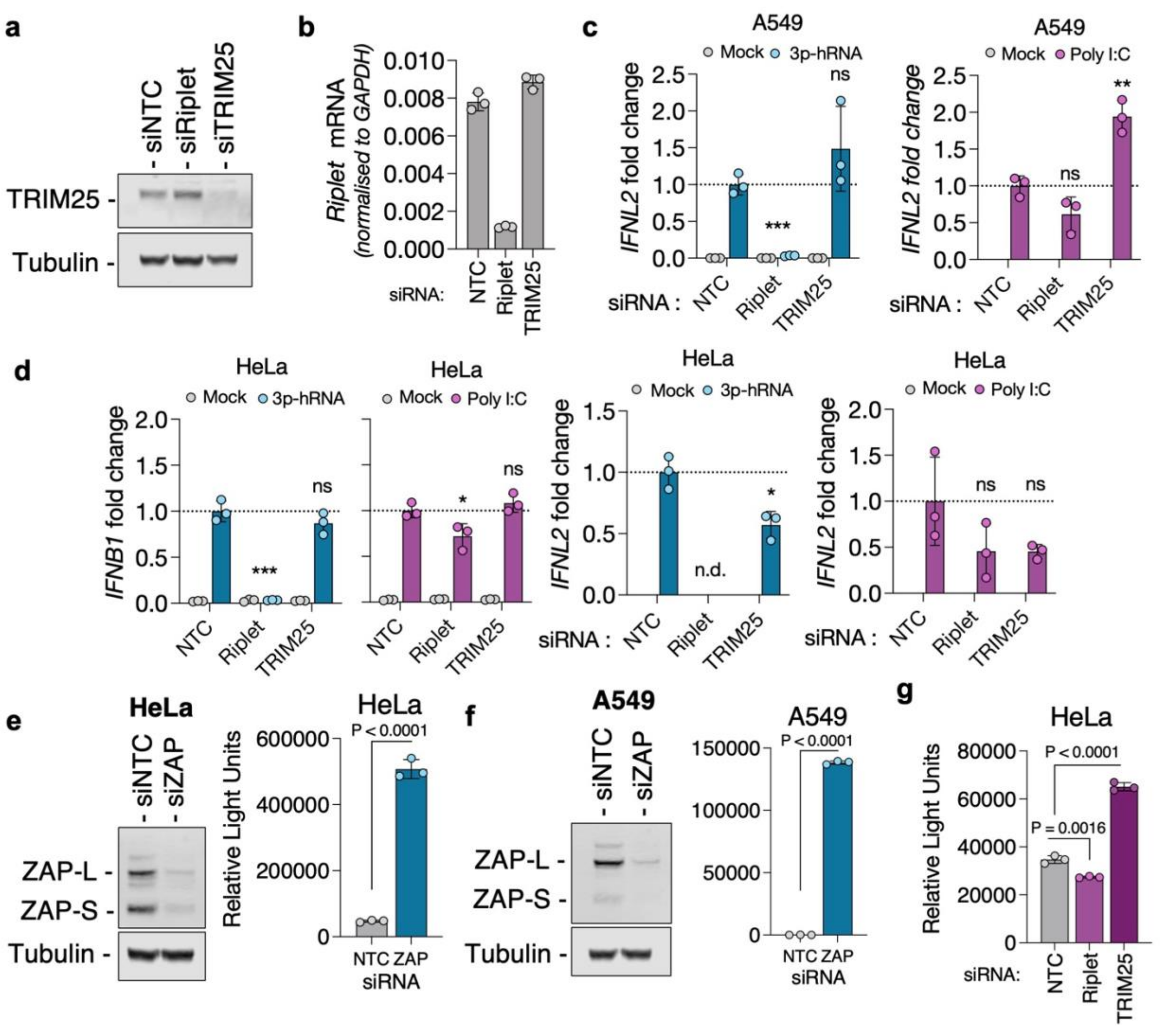


**Supplementary Figure 3. TRIM25 and Riplet support ZAP and RIG-I activity, respectively**. TRIM25 and Riplet were depleted from A549 cells using siRNAs and 48h later cells were analyzed by western blotting (a) and qPCR (b). Riplet and TRIM25 were depleted from A549 cells using siRNAs and 48h later cells were transfected with 3p-hRNA or poly I:C. After 8h after transfections, RNA was extracted and IFNL2 was quantified by qPCR (c). Riplet and TRIM25 were knocked-down from HeLa cells using siRNAs; cells were then transfected with 3p-hRNA or poly I:C and, after 8h, RNA was extracted and *IFNB* and *IFNL2* transcripts were quantified by qPCR (d). ZAP was depleted from HeLa cells (e) or A549 cells (f) using siRNAs. After 48h, cells were analyzed by western blotting or infected with a ZAP-sensitive EV71. Luciferase activity was measured 24h later. Riplet- or TRIM25-depleted HeLa cells were infected with ZAP-sensitive nLuc-EV71, and 24h later luciferase activity was quantified (g).

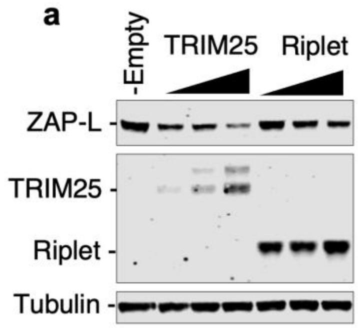


**Supplementary Figure 4. TRIM25 expression impacts ZAP protein levels.** HEK293T TRIM25$^{-/-}$ ZAP$^{-/-}$ were transfected with increasing amounts of plasmids encoding TRIM25-HA or Riplet-HA and ZAP-L expression was analyzed by western blotting (a).

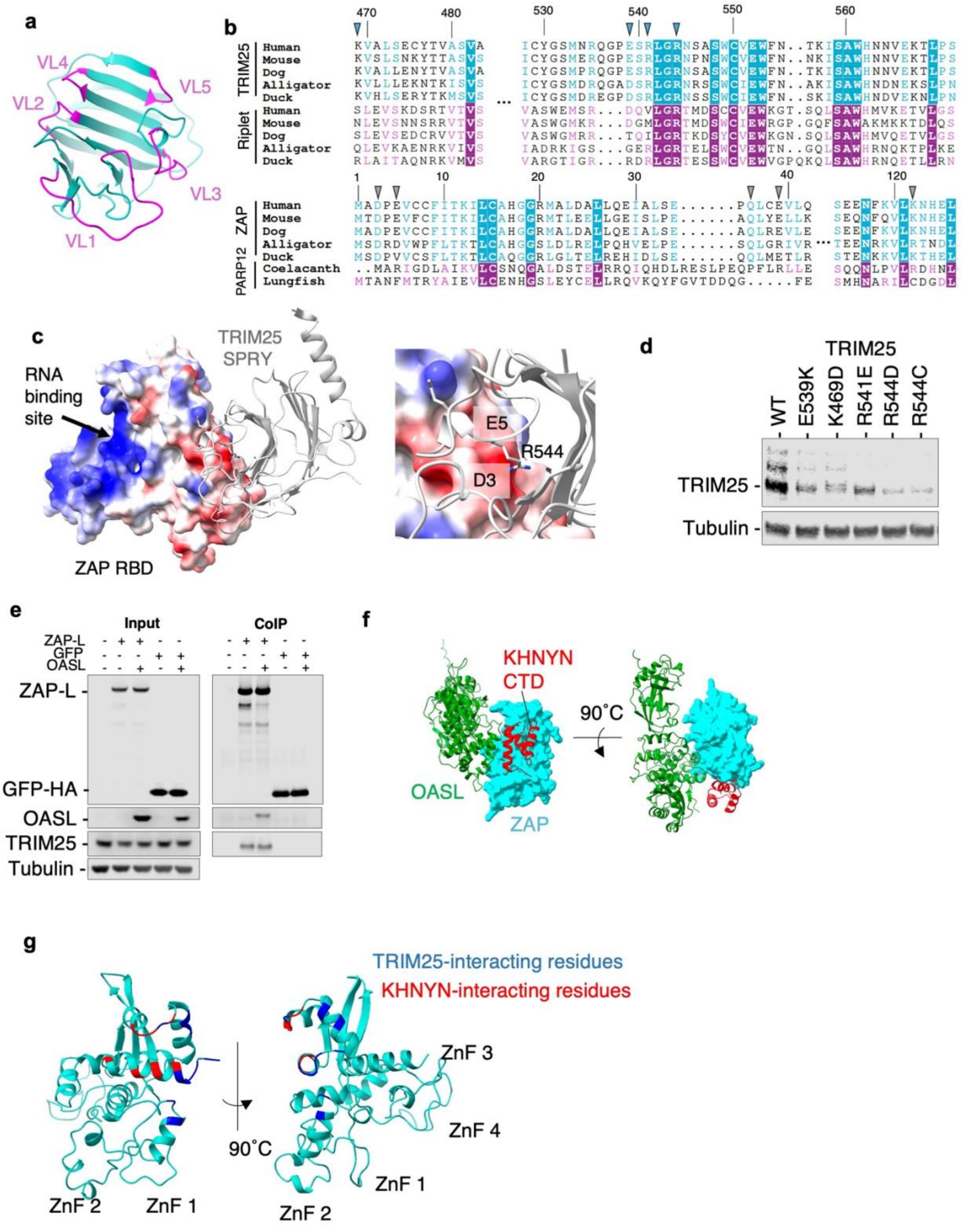


**Supplementary Figure 5.** Structure of the SPRY domain of TRIM25 with variable loops highlighted in purple (a). Multiple sequence alignment of orthologues of TRIM25, Riplet, ZAP and PARP12 (b). Predicted structure of ZAP-TRIM25 showing electrostatic surface (c). Inset showed positively charged R544 residue of TRIM25 accommodated by a negatively charged pocket formed by residues D3 and E5 of ZAP (c). Expression levels of single mutants of TRIM25 as indicated (d). HEK293T were transfected with plasmids encoding ZAP-L-HA, GFP-HA and OASL; cell lysates were used to purified ZAP- or GFP-complexes using an anti-HA antibody and analyzed by western blotting (e). Structure of the RNA-binding domain of ZAP (in blue) and the C-terminal domain of KHNYN (red, KHNYN CTD, obtained from PDB:9BGL) and the predicted sturcture of OASL (in green) (f). Predicted structured of the RNA-binding domain of ZAP (light blue) with residues known to interact with KHNYN in red (described by Bohn et al. [24] ) and residues predicted to interact with TRIM25 in blue (g).
